# Exploring genes of rectal cancer for new treatments based on protein interaction network

**DOI:** 10.1101/037531

**Authors:** Wenjing Teng, Chao Zhou, Yan Li

## Abstract

**Objective:** To develop a protein-protein interaction network of rectal cancer, which is based on genetic genes as well as to predict biological pathways underlying the molecular complexes in the network. In order to analyze and summarize genetic markers related to diagnosis and prognosis of rectal cancer.

**Methods:** the genes expression profile was downloaded from OMIM (Online Mendelian Inheritance in Man) **database**; the protein-protein interaction network of rectal cancer was established by Cytoscape; the molecular complexes in the network were detected by Clusterviz plugin and the pathways enrichment of molecular complexes were performed by DAVID online and Bingo (The Biological Networks Gene Ontology tool).

**Results and Discussion:** A total of 127 rectal cancer genes were **identified to differentially express** in OMIM Database. The protein-protein interaction network of rectal cancer was contained 966 nodes (proteins), 3377 edges (interactive relationships) and 7 molecular complexes (score>7.0). **Regulatory effects of genes and proteins were focused on cell cycle, transcription regulation and cellular protein metabolic process**. Genes of *DDK1, sparcl1, wisp2, cux1, pabpc1, ptk2* and *htral* **were significant nodes in PPI network**. The discovery of featured genes which were probably related to rectal cancer, has a great significance on studying mechanism, distinguishing normal and cancer tissues, and exploring new treatments for rectal cancer.

---

Rectal cancer is one of the common fatal malignant tumors in the world, which is secondly lethal cancer in United States as well as thirdly in Europe. The research has improved systemic treatment, such as micro diagnosis. However, despite advances in detection and care, morbidity and mortality from rectal cancer continues to be high[1].Early detection and diagnosis can be great significance to reduce mortality and improve prognosis, as well as identify those who were at the highest risk duo to improving triage for treatment has the greatest impact on rectal cancers treatment. Studies shown that rectal cancer has a prime paradigm for cancer genetics, which can be prevention by early detection of the pre-disease (neoplastic) state. Therefore, the roles of genetics in rectal cancer have became critical to the missions of disease prevention, early detection and effective treatment

As report goes, approximately 10% of well defined hereditary rectal cancer has syndromes[2]. Hereditary rectal has some benefits, such as many of it are growing slowly, family members are armed with the knowledge of potential risk of associated carcinomas and empowerment to reduce the disease burden.

A large number of mutation, mismatch, inactivation of tumor suppressor genes and activations of oncogenes are involved in the process of rectal cancer[4].Up to now, genetic marker of rectal cancer can be summarized by the following six aspects genomic instability, CpG island methylator phenotype, specific microRNAs, histone modification, gene mutation and protein biomarkers. The protein-protein interaction network which is a model of biological molecular interactions can more clearly show the genes, proteins and pathogenesis in the development process of the disease[5]. These genetic traits may partially explain the geographical variance in rectal cancer incidence and mortality as well as the differences between hereditary and sporadic rectal cancer [6].

## 1 Materials and Methods

**Design: Study on enrichment of genomic biological pathways.**

**Method:**

**Data acquisition:** OMIM(Online Mendelian Inheritance in Man) is a comprehensive,authoritative, daily updated human phenotype database, containing more than 12000 genes of all human genetic diseases, and mainly focusing on hereditary diseases. In addition, text messages, related reference information, sequence records, maps, and related databases are available for each gene[7-8]. This study had started from August 22(2015), searched "rectal cancer"in the OMIM database and obtained human genes associated with rectal cancer information.

**The construction of gene/protein interaction networks:** rectal cancer associated genes were submitted to Cytoscape 3.2.1 plug-in Agilent Literature Search 2.7.7 (USA Agilent Technologies company) and Pubmed [9]. False positive interaction information was removed from retrieval results. Then, gene/protein interaction relations were read in Cytoscape 2.8.2 and visualized [10].

**Network analysis:** MODE algorithm in Cytoscape 3.2.1 web analytics plug-in Clusterviz of 1.2 was administrated to make the correlation analysis for the area of the construction of biological networks [11-12]. By analyzing the network structure, proteins were grouped to form molecular compounds in the entire network and were shown in Cytoscape according to the correlation integral value. The areas with integral value higher than 3 were regarded as molecular compounds. The gene/protein names contained in the molecular compounds were submitted to The Database for Annotation, Visualization and Integrated Discovery. [13-14] By retrieving Kyoko Encyclopedia of Genes and Genomes (KEGG) Database, biological pathways involved in chronic myelogenous leukaemia heredity were identified. Then the Biological pathways data were submitted to **Bingo** (in Networks Gene Ontology tool) for enrichment analysis.

**Main outcome measures:** Protein networks were constructed based on the rectal cancer-related genes, nodes (proteins) and edges (interaction between), molecular complexes in the network and its associated interaction points and nodes (protein) and the edge (interaction between), analyze the biological pathways has involved in the molecular complexes.

## 2 Results

### 2.1 Rectal **cancer related genes in OMIM**

According the OMIM database retrieval, it can found that 127 genes were reported to be associated with rectal cancer. After screening and deleting duplicate genes, 127 related genes were identified, which shown in **Table1**.

### 2.2 Protein interaction networks

Through text mining,127 genetic-related genes shown that there was a network diagram with 638 nodes (proteins) and 1830 edges. As shown in **Figure 1-3**, the diamond is represented OMIM genetic disease related proteins, while the round represented the proteins obtained from text mining

**Figure 1.**
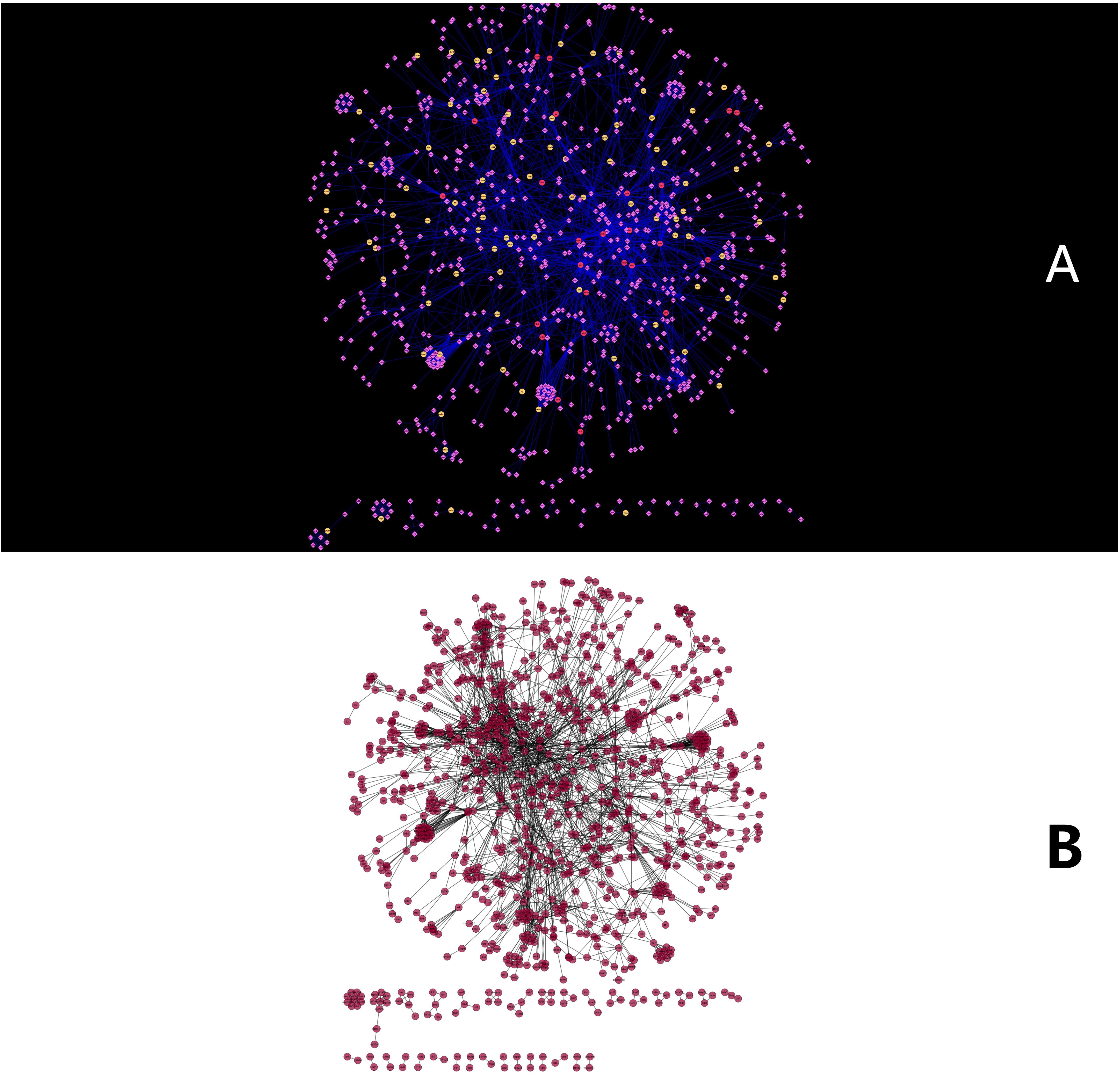
(A+B). Network map of protein interaction (Overall)

**As shown in figure 1**, the protein-protein interaction network underlining rectal cancer is extremely complex. The edges intersect with each other and several clusters emerge in the figure. The more edges among the genes, the correlation of the genes is more tightening. So the genes formed the round network at the top of figure1 connected with each other more tightly than the genes in the bottom.

After removing the protein molecule in the network, the relationship between proteins in network became more clear, in the centre part of protein-protein network, the relationship between protein complexes emerged is closer, however, at the farther edge of the network, the relationship of it became looser, as shown in **Figure2**.

**Figure 2.**
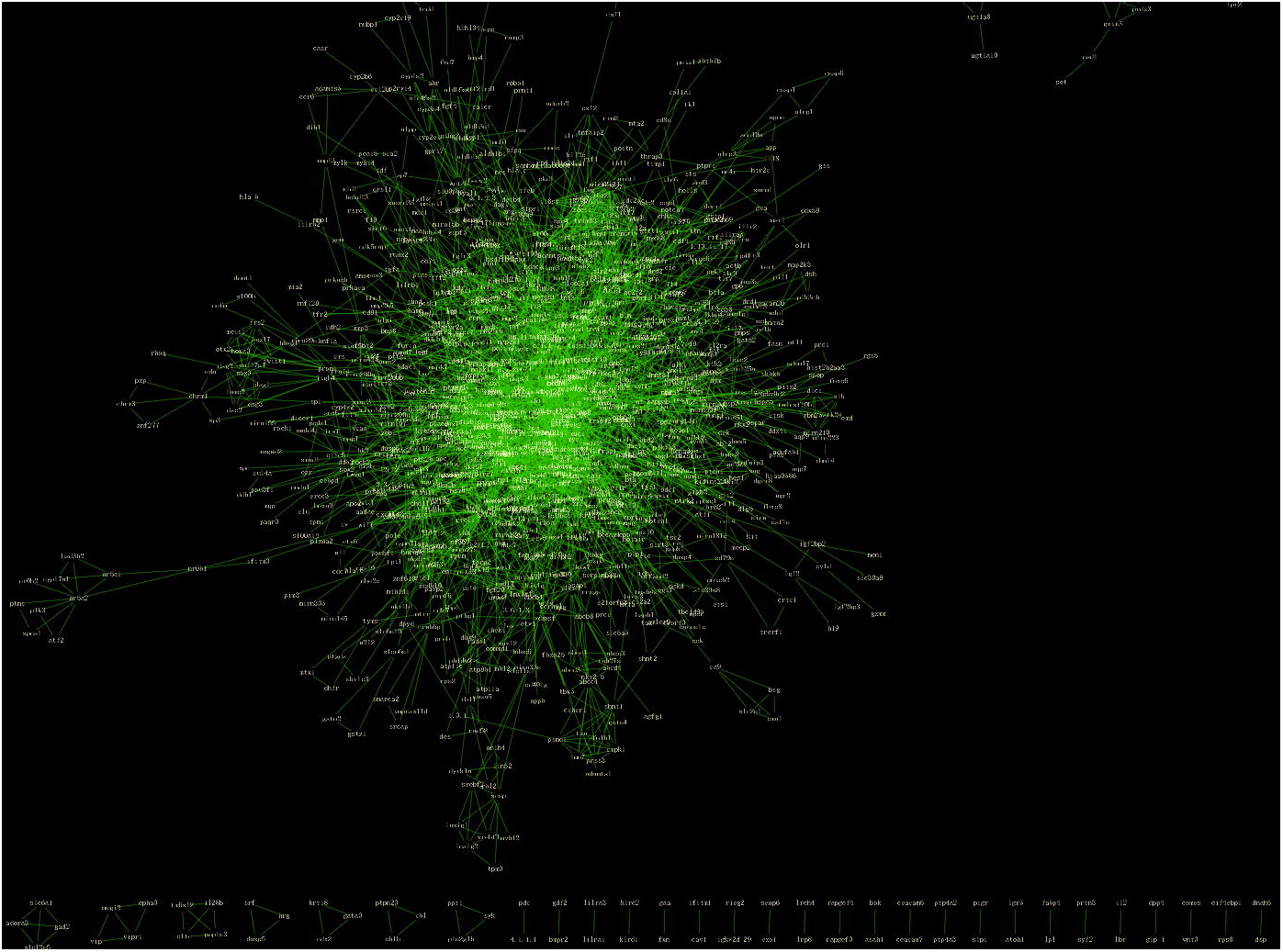
the relationship of proteins

AS shown in **Figure3**, it can be found some common genes and pathways such as jak2-stat1,Kras, P53,etc as well as genes associated with them.

**Figure 3.**
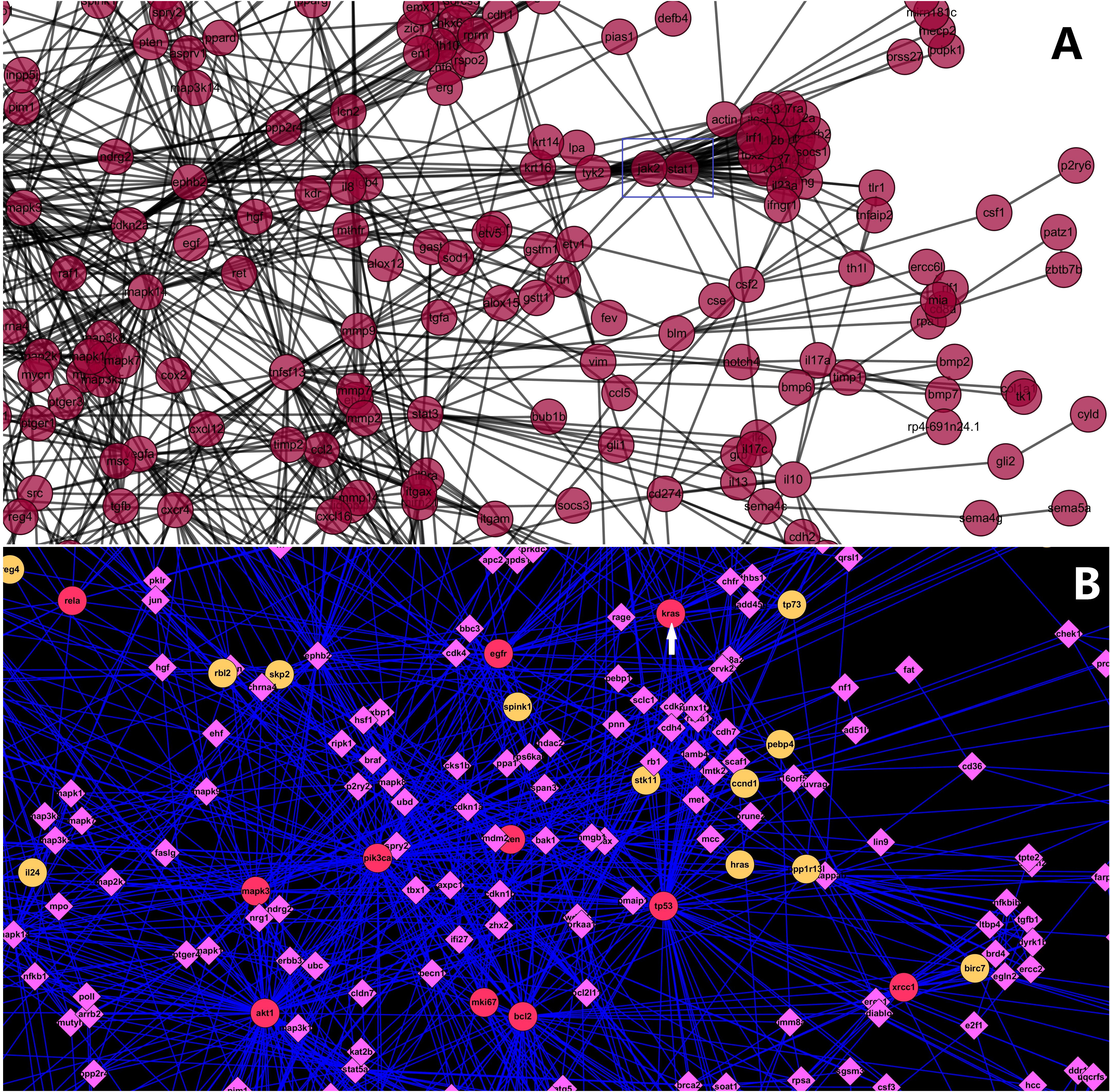
pathway jak2-stat1 relationship of gene *Kras*

### 2.3 Network topology attribute analysis

Network topology attribute analysis shown that the connectivity of nodes in the network (the number of nodes in the network) obeys descending distribution, i.e. with the increase of edges connected to the node, correspondingly the number of nodes decrease, so it can be seen that the gene - protein interaction networks are scale-free networks [15]. We found that the connectivity of nodes in the network greater than or equal to 25 corresponds to a sharp reduction in the number of nodes **Figure 4**. Therefore, we regarded the nodes which the connectivity is greater than / equal to 20 as the key nodes (hub). Key nodes (connectivity score)included: hif1a(25), cdkn1b(25), rb1(26), plau(26), brca1(27), asns(28), pcna(28),Vegfa(29), stat1(29), cdkn2a(29), egfr(30), tp53(71).

**Figure 4.**
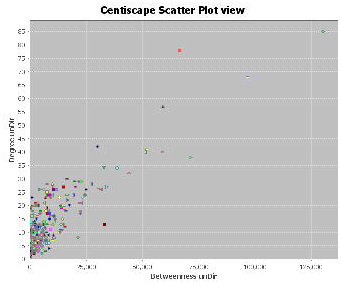
Connectivity degree of each node and betweenness Comparison

(The horizontal axis represents between ness, and the ordinate represents the connectivity degree of protein interaction network. And the graphic in the table represents each node in the network). It can be seen that the connectivity (the number of nodes in the network) of nodes in the network obeys descending distribution, while the connectivity is greater than / equal to 25, the number of nodes corresponding to a sharp decrease.

### 2.4 The detection of molecular complexes

According MCOMD algorithm analysis, analyse the correlation between genes in network and calculate the score, the number of nodes, the number of edges. There is a total of 74 molecular complexes and 7 of them showed correlation integral values higher than7.0.

**Figure 5.**
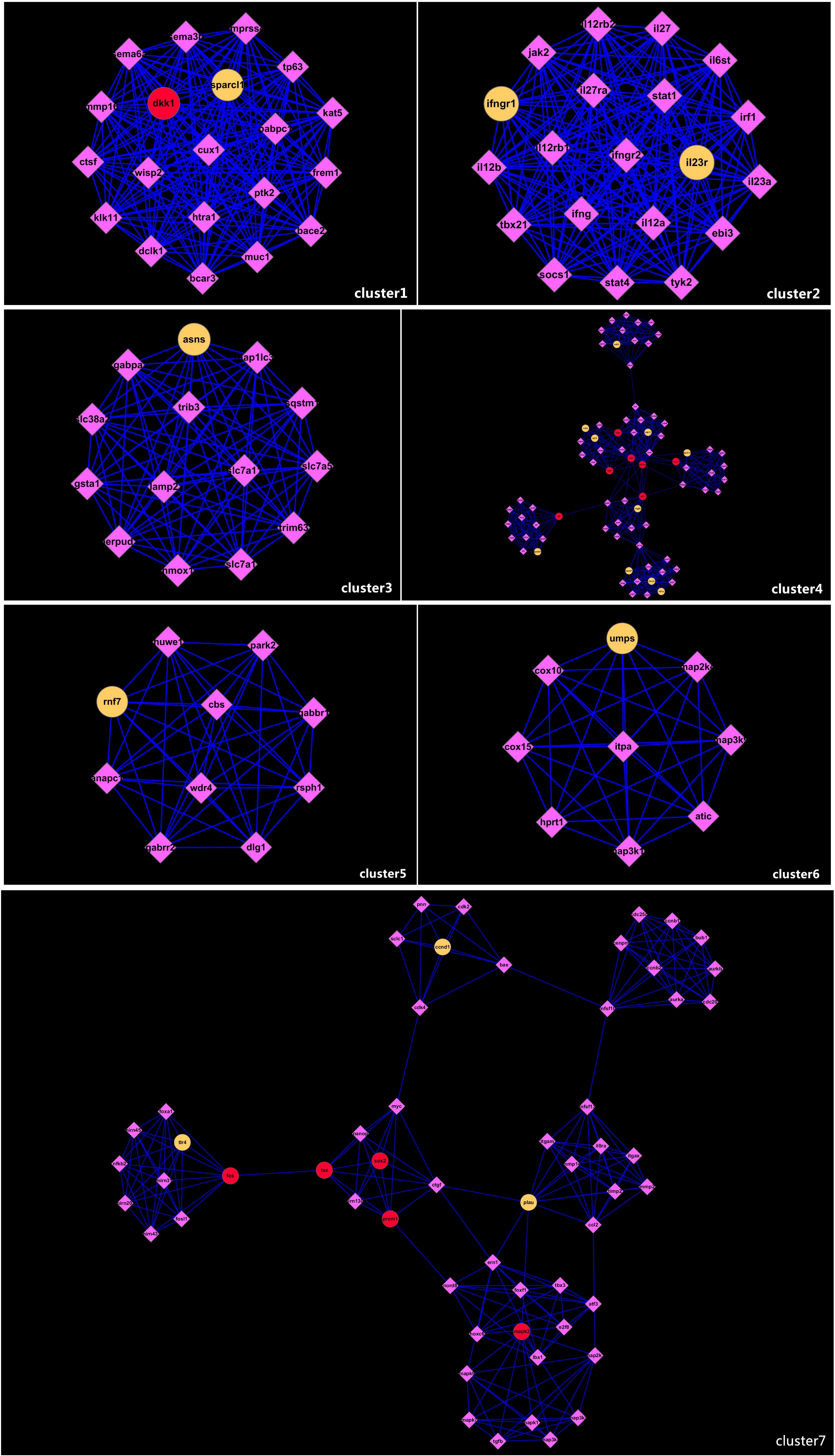
Molecular complexes obtained by MCOMD algorithm analysis. **Rank1** score20 Node20 Edges190 **Rank2** score20 Node20 Edges190 **Rank3** score14 Node14 Edges91 **Rank4** score11.662 Node78 Edges449 **Rank5** score3.143 Node21 Edges66 **Rank6** score9 Node9 Edges36 **Rank7** score7.673 Node56 Edges221

The details of the relationship of proteins in molecular complexes 7 was shown in Figure 6

**Figure 6.**
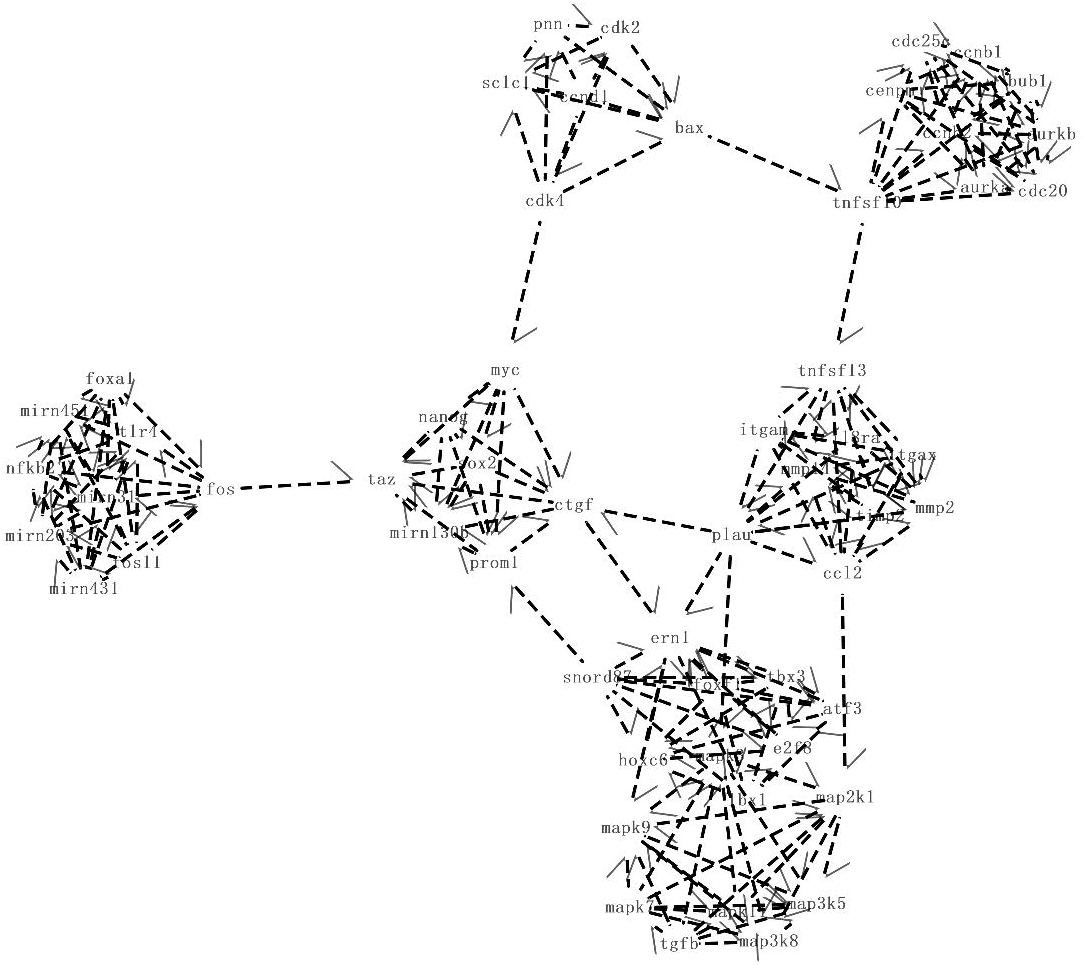
the relationship of proteins in molecular complexes 7

The whole network has a total of six parts, including mitogen-activated protein kinase, Specific microRNAs, SRY-related HMG-box, matrix metalloproteinases, cyclinD.

### 2.5 **Molecular complex pathway** enrichment

Submit the 7 names of protein molecule complexes online to obtain the relevant pathways in **Table 2**. The Protein molecule biological pathways of complexes 1 are not exist,and other complexes contain different pathways.Regulatory effects of genes and proteins mainly focused on cell cycle, transcription regulation, and cellular protein metabolic process.

**Table 2.**
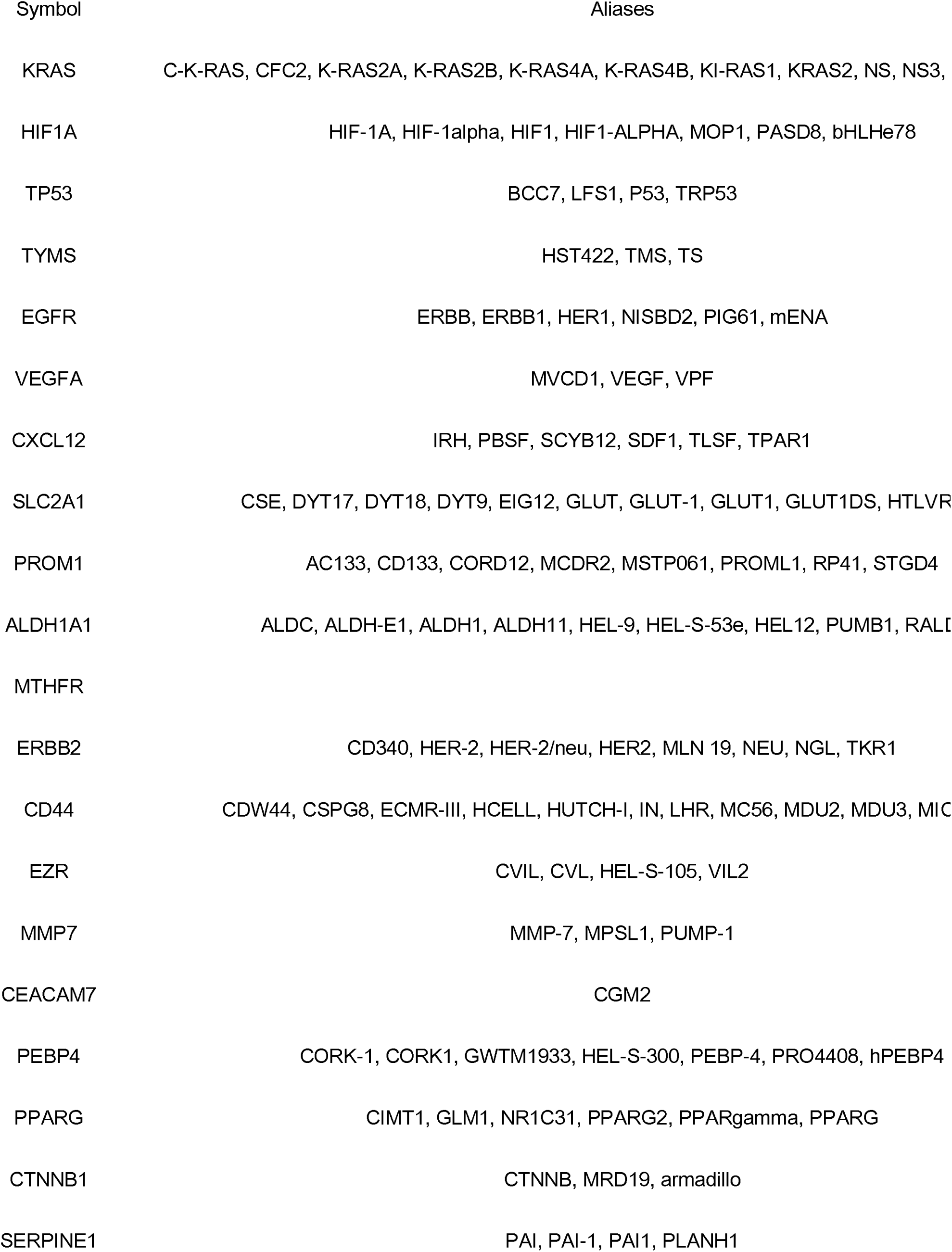
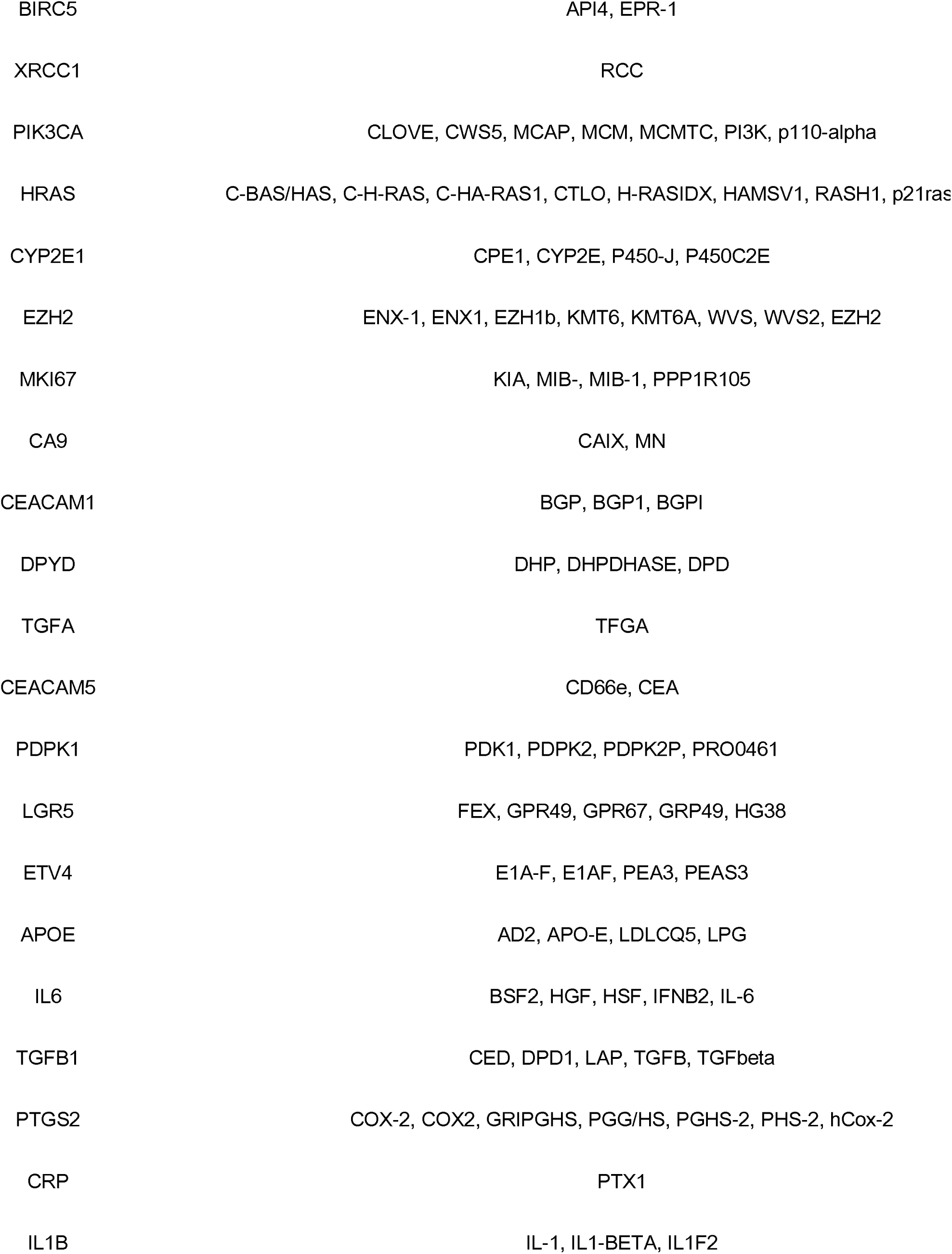
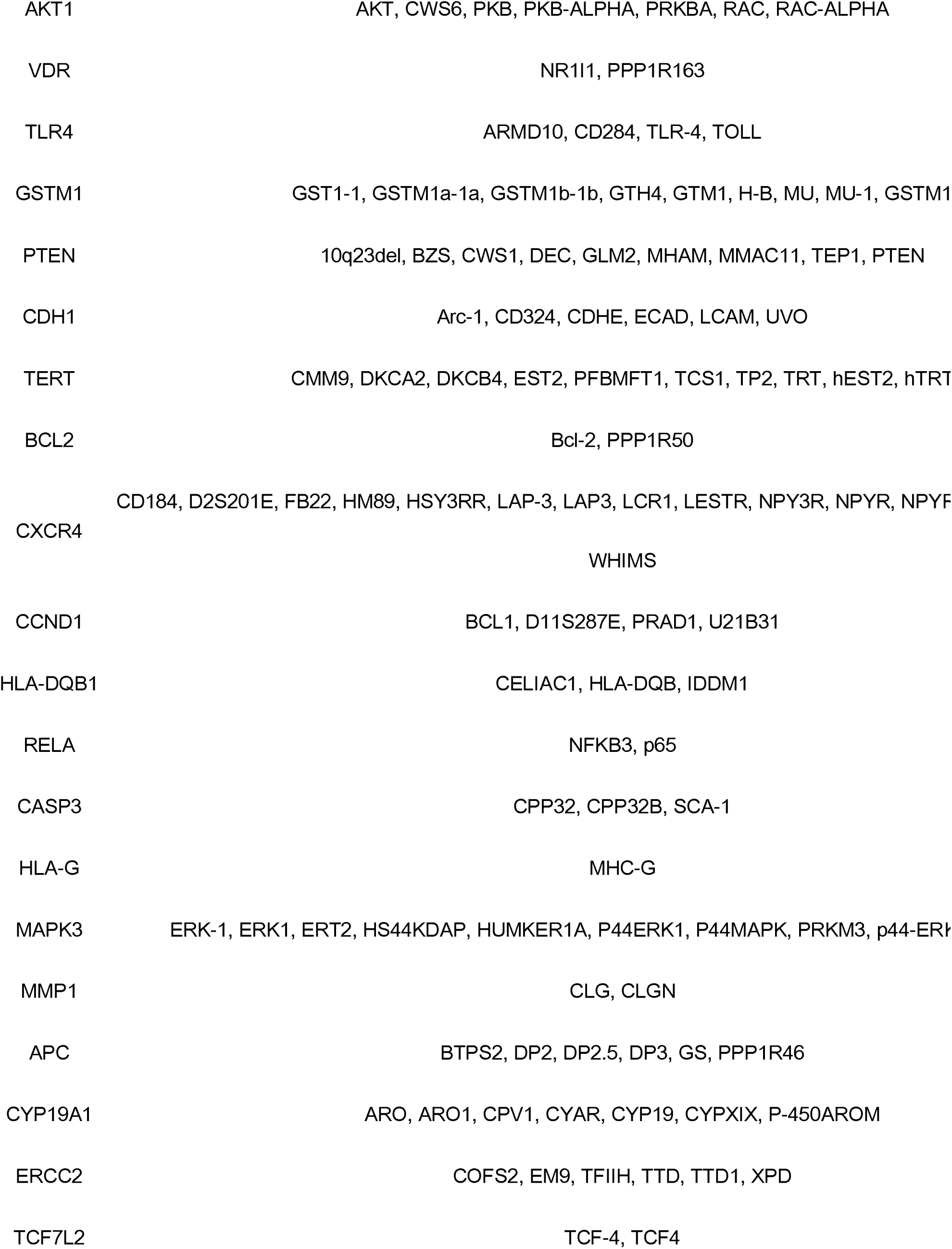
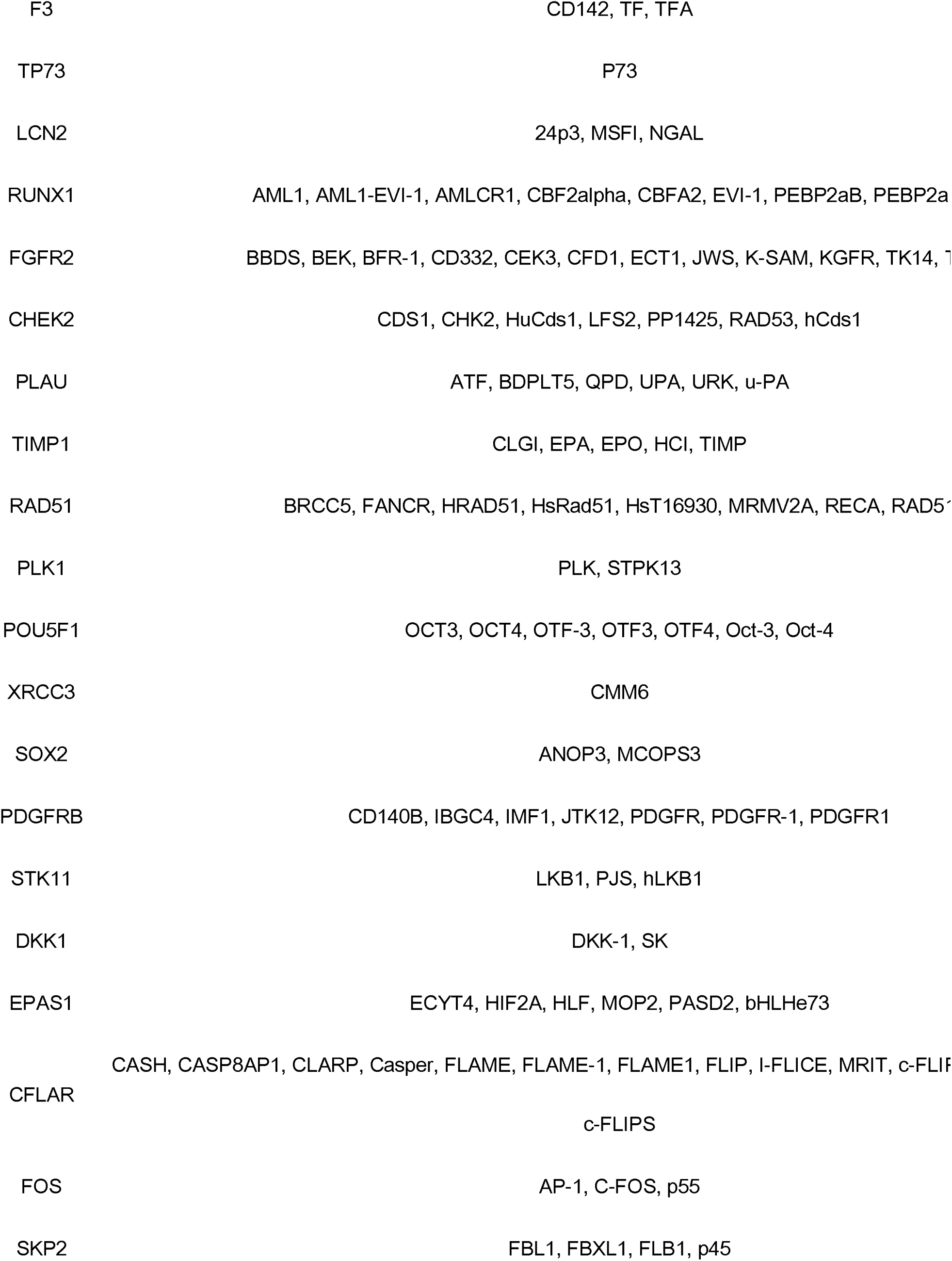
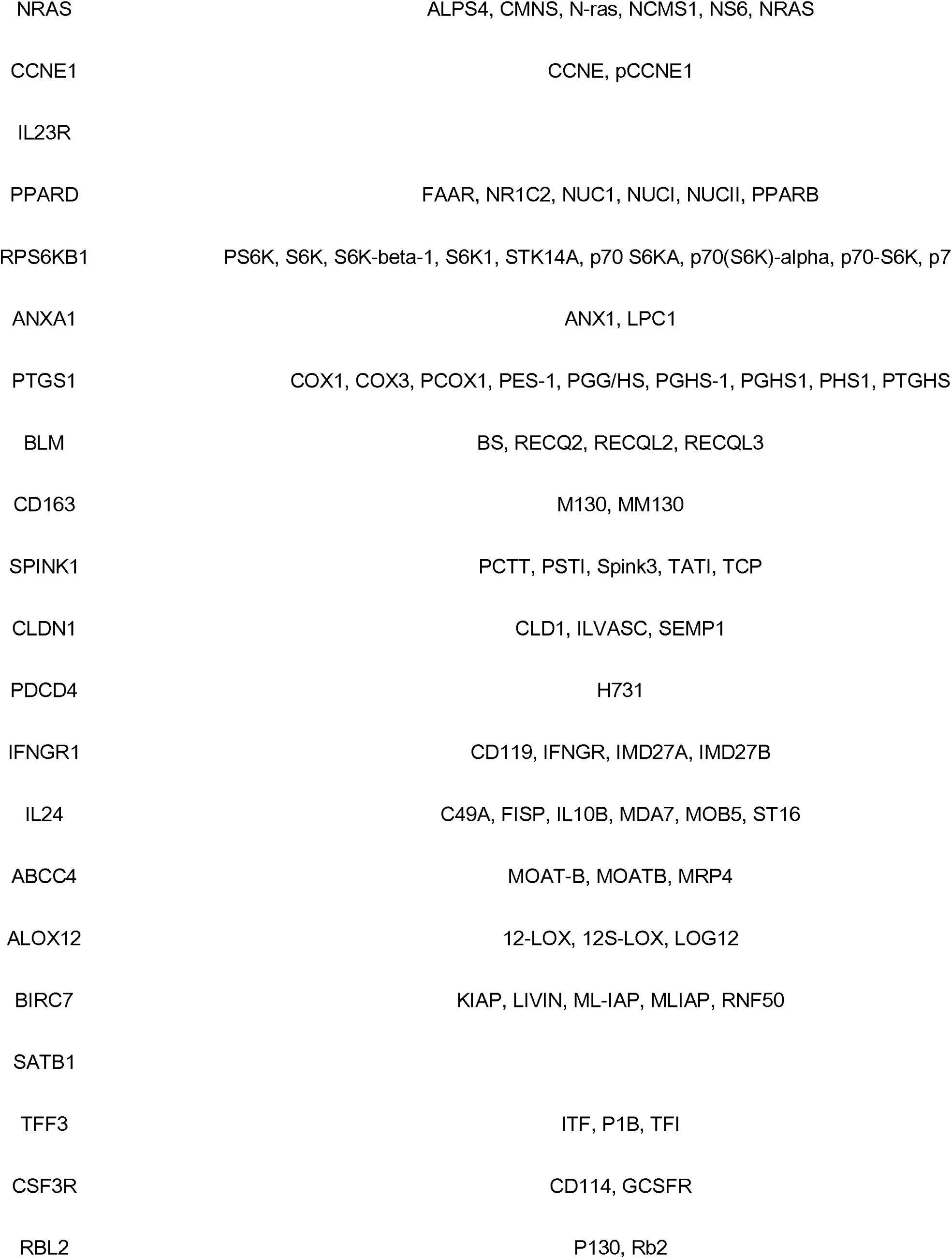
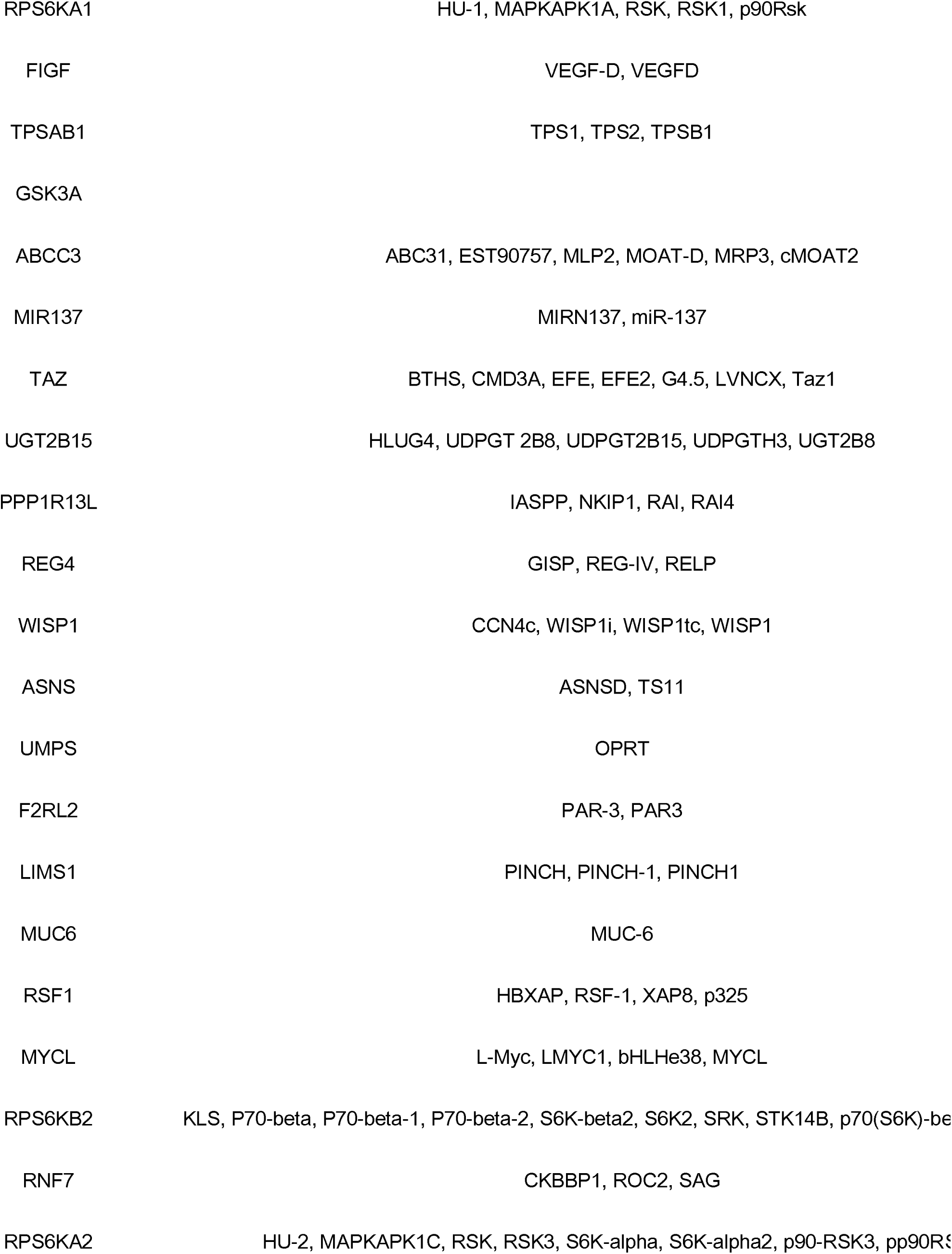
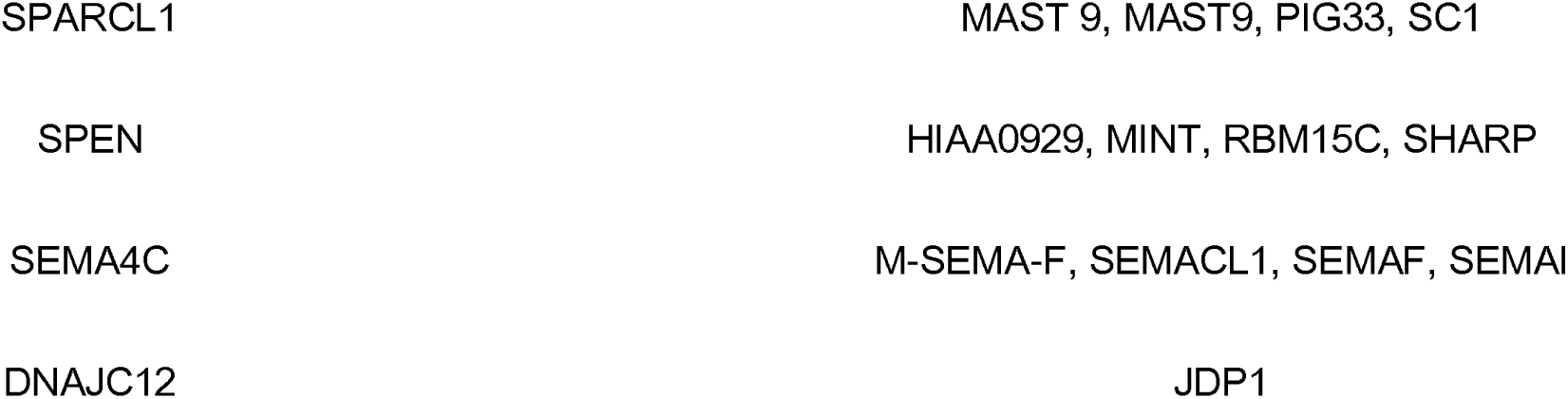
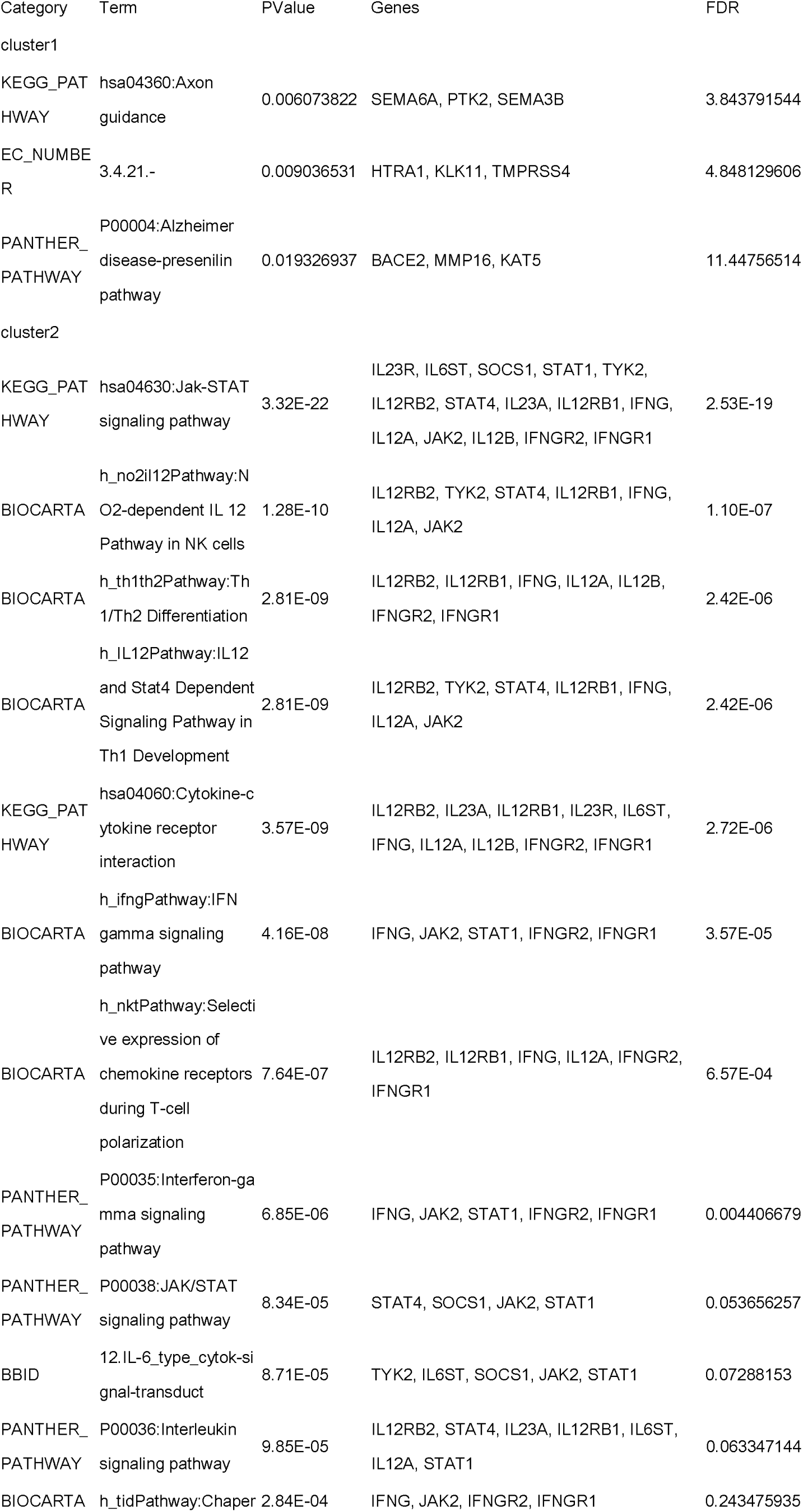
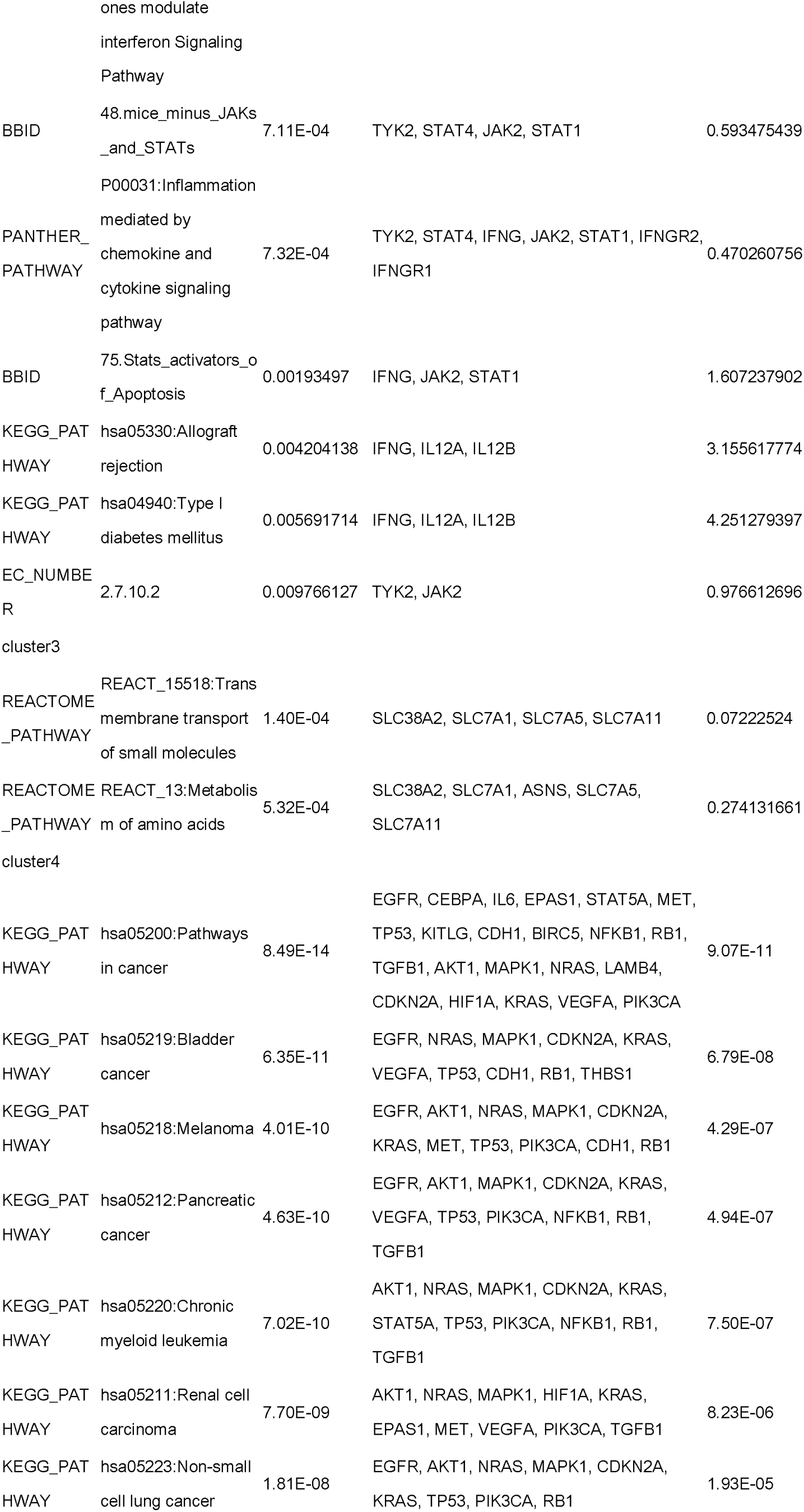
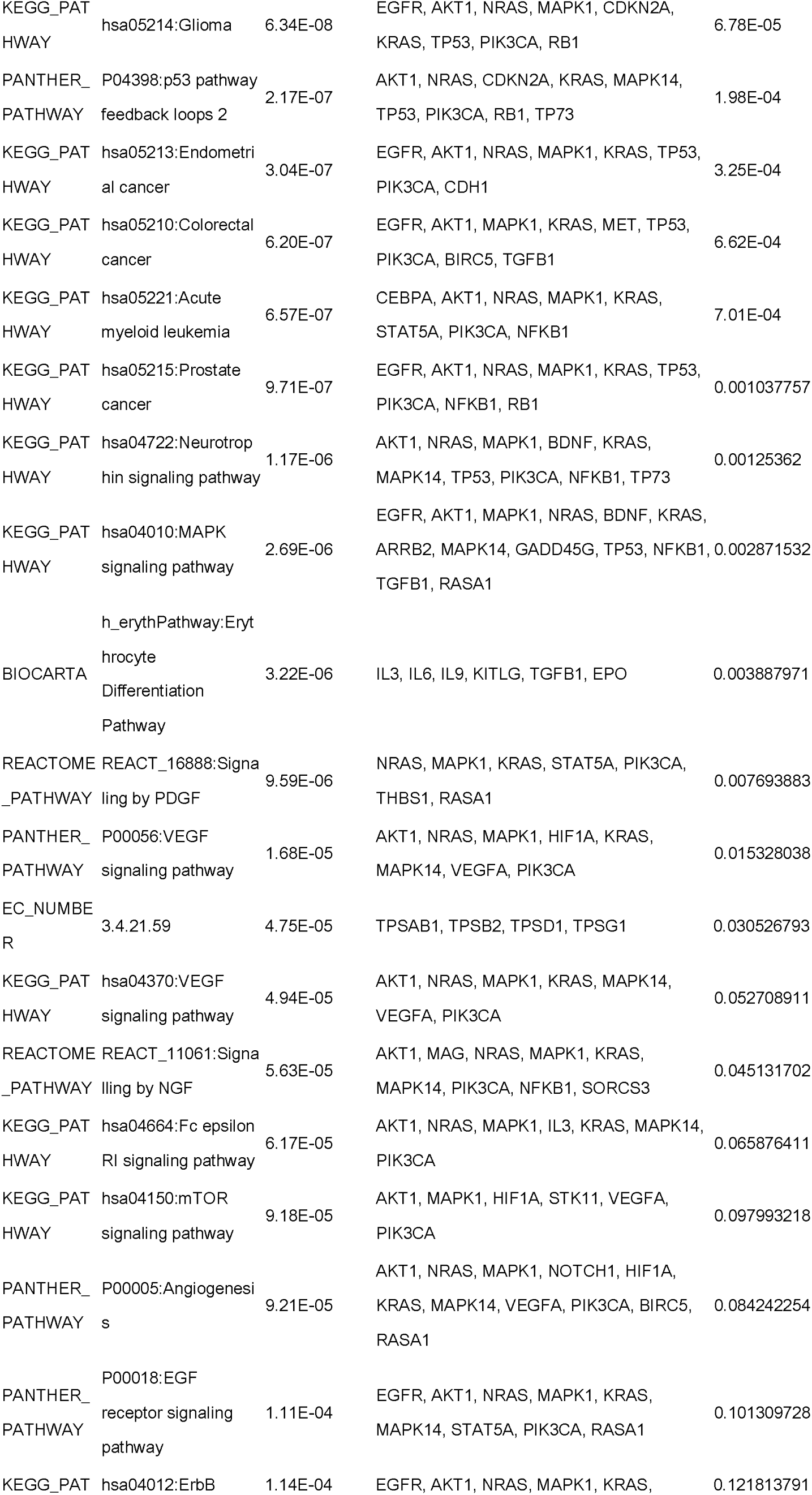
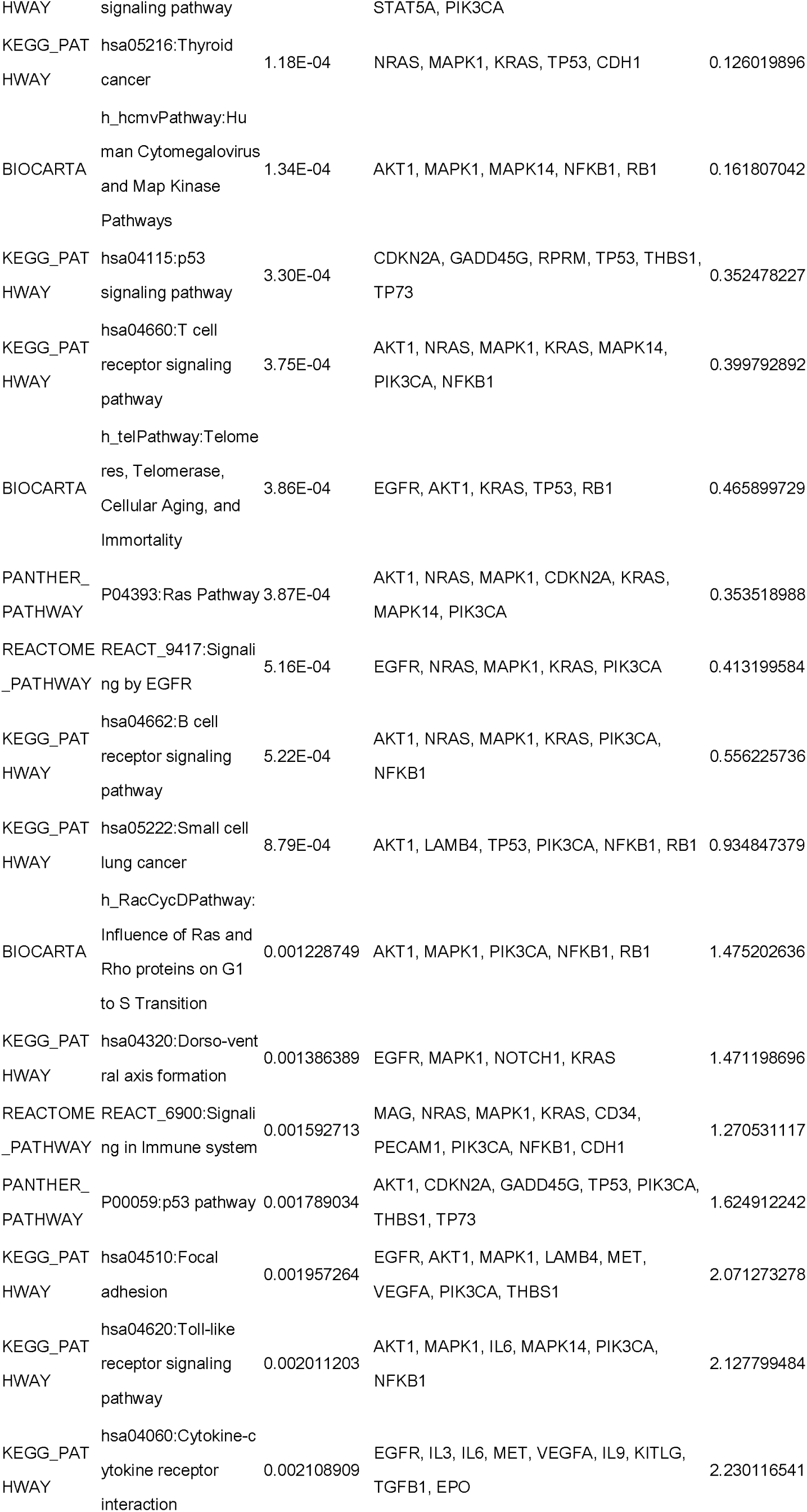
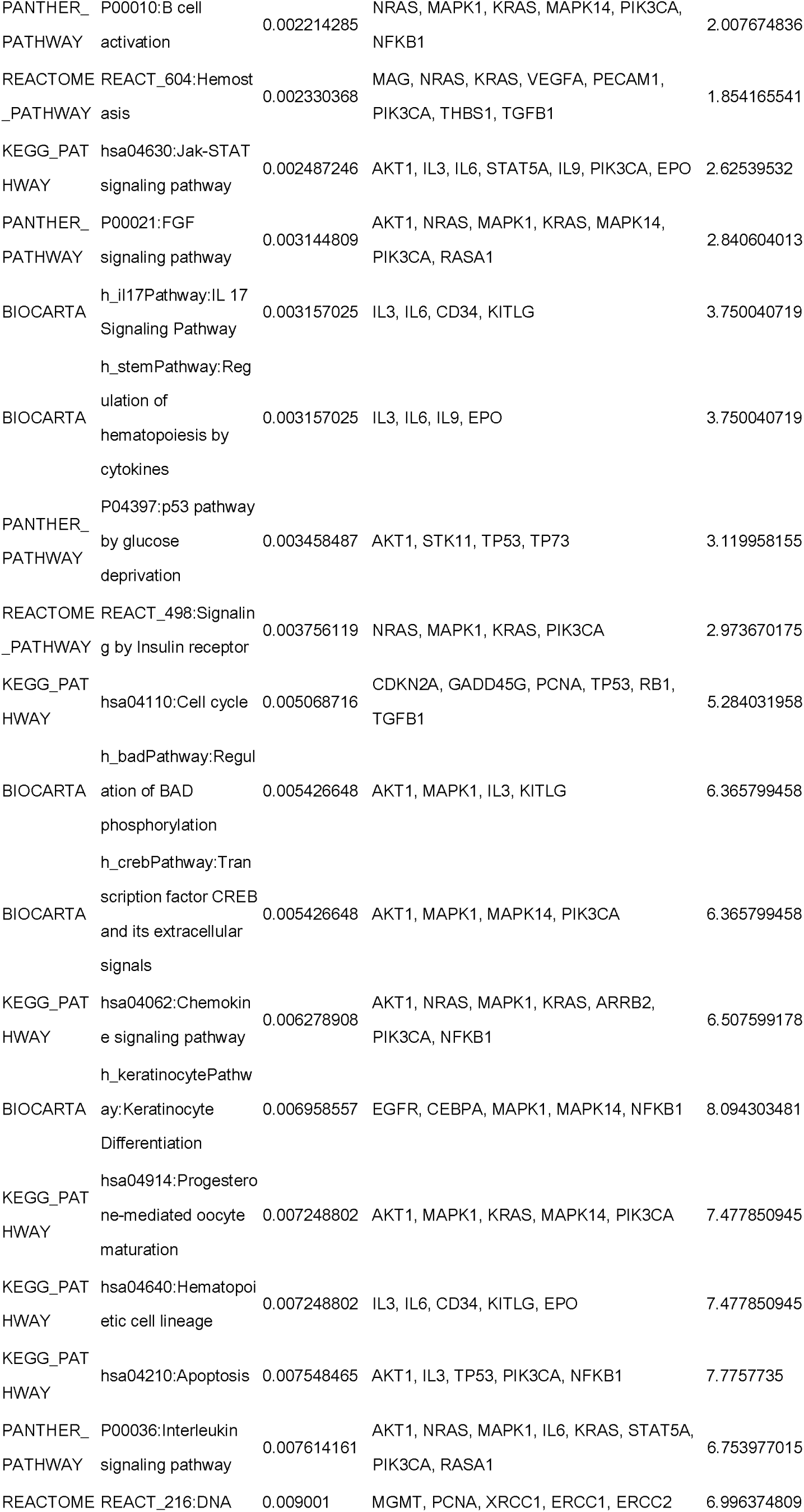
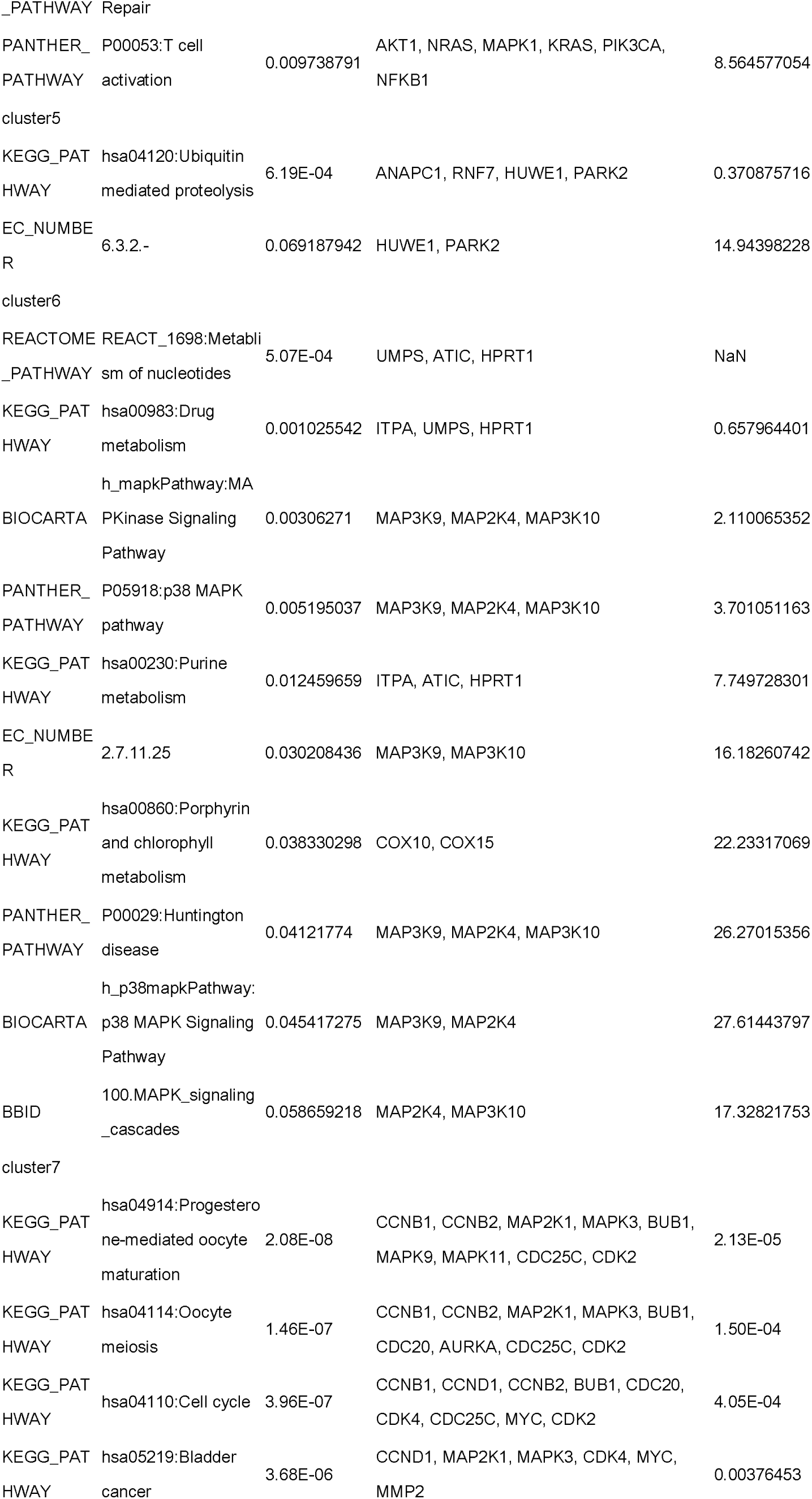
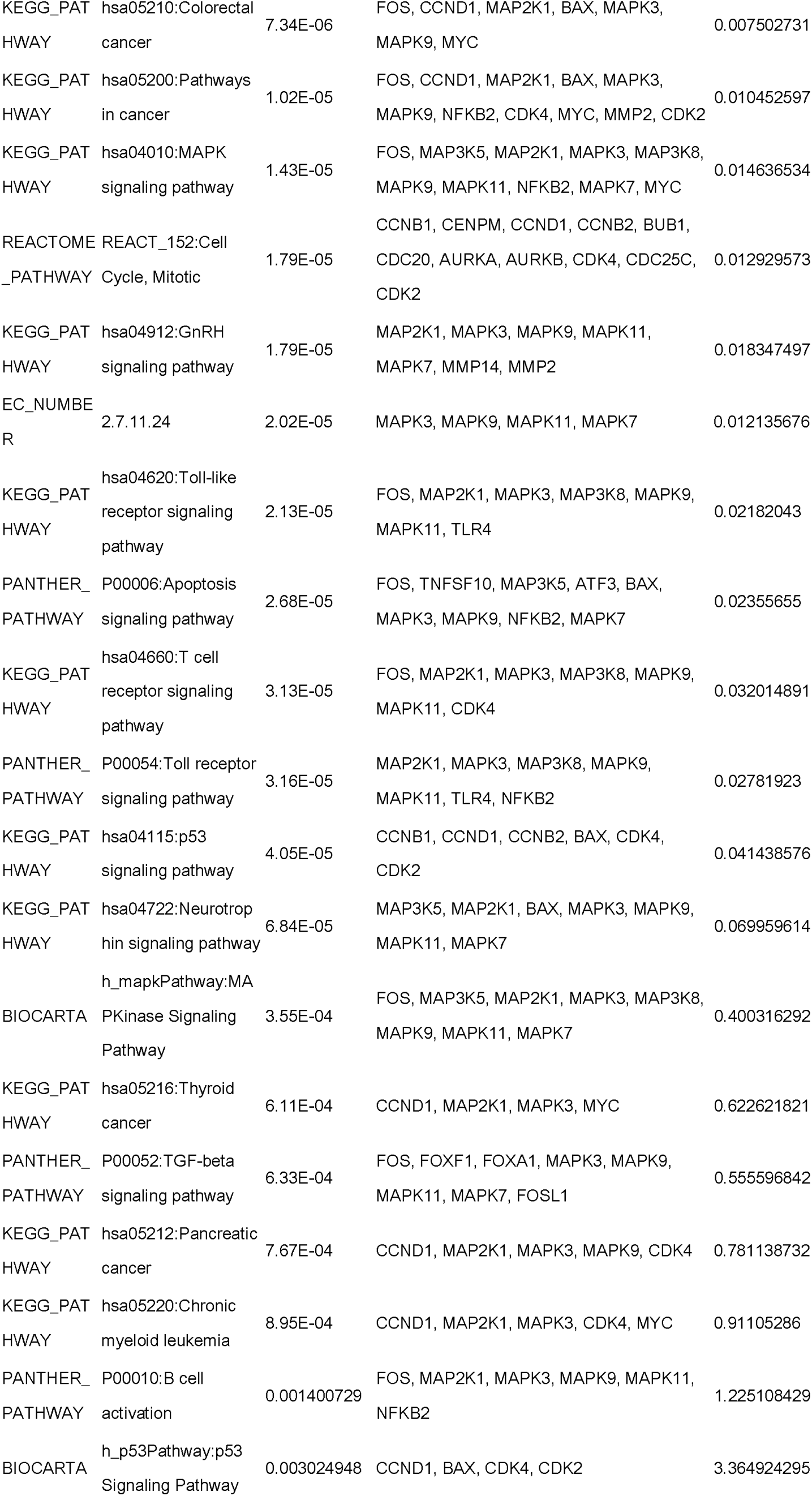
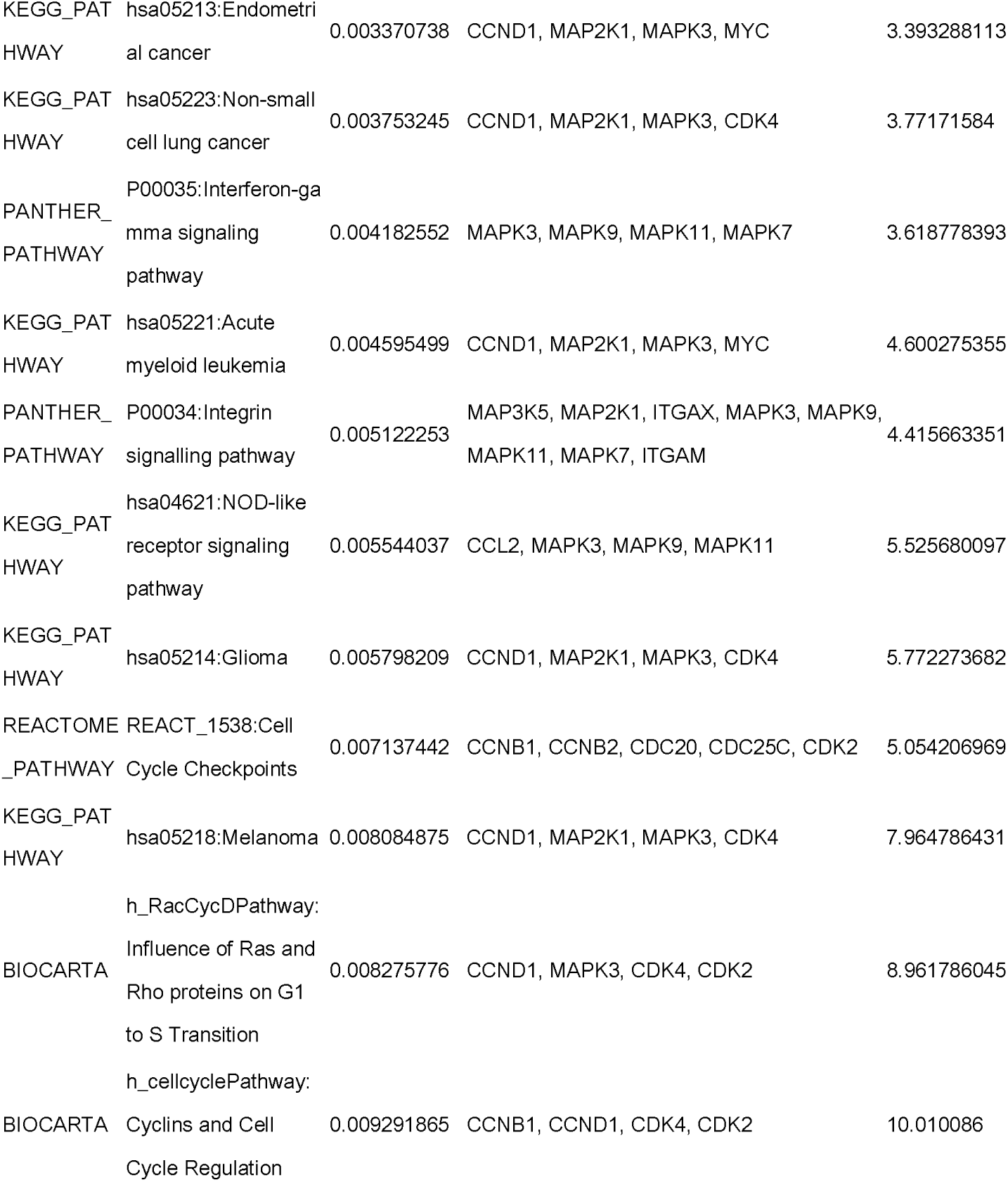
Enrichment of pathways related to molecular complexes

Binggo results show the gene oncology hierarchical network to the biological processes, **Figure 7**. The size of the node represents the number of the genes, the depth of the node color represents the P values.Diagrams have presented the main biological processes of cluster2,3,5,containing metabolic regulation, transcriptional regulation, biosynthesis, cell differentiation and gene expression regulation and signal transduction, etc.

**Figure 7.**
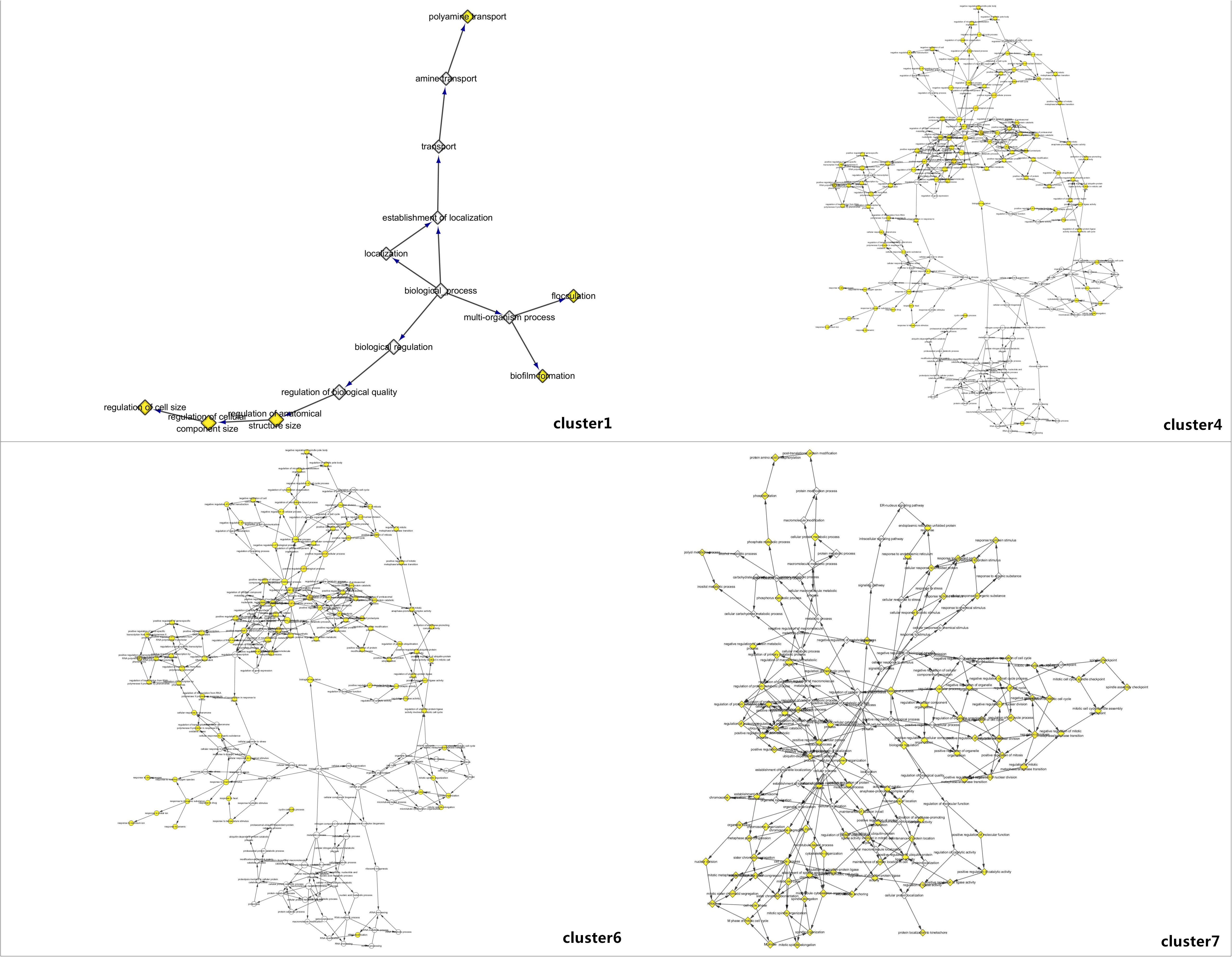
**A Bingo results show that the basic structure of the biological cluster1** **B Bingo results show that the basic structure of the biological cluster4** **C Bingo results show that the basic structure of the biological cluster6** **C Bingo results show that the basic structure of the biological cluster7**

## 3 Discussions

Based on a large number of research references, the progression of rectal cancer is a multistep process, containing overcome apoptosis, inhibiting senescence (infinite proliferation), secretion proliferation signal itself, not sensitive to growth signals, angiogenesis and invasion, which requires a large number of genes and proteins in action[16].Genetic markers of rectal cancer can be summarized from the following six aspects: he first part is genomic instability, close to 12-17% of sporadic rectal cancer cases exist with microsatellite instability (MSI). At present, microsatellite instability - high - H (MSI) has become a positive prognosis of rectal cancer patients’ overall survival[17]. 50-85% of Rectal cancer patients has chromosome aberration frequency, chromosome instability (CIN +) usually be associated with colorectal cancer patients’ overall survival, progression-free survival (PFS) and poor prognosis after 5 - fluorouracil therapy; the second part is CpG island methylator phenotype. Nearly 29.6% of the rectal cancer patients shown CpG island methylation phenotype - high (CIMP - H) [18], however, its value is still in research. Specific microRNAs and histone modification are another two aspects, which mainly associate with colorectal cancer patients’ overall survival and progression-free serial, tumor metastasis, local invasion, tumor volume, tumor staging, treatment results, relapse and drug resistance.

What is more, gene mutations and protein biomarkers have brought a special significance. The APC gene mutant p.D1822V containing homozygous V/V would reduce the risk of rectal cancer[19]. According to the APC (rs565453 y rs1816769) and CTNNB1 (rs229303) gene polymorphisms, the death risk stratification in patients with rectal cancer can be analysed. The APC gene mutant p.I1307K as risk factors of the rectal cancer among Ashkenazi Jewishes has been found[20]. The loss of PTEN gene in 22% of rectal cancer patients has caused no response to the EGFR inhibitors and higher risk of death [21]. And the clinical application of protein molecular markers colorectal cancer mainly included: early diagnosis(hnRNP A1,kininogen-1,adipophilin, Apo AI and C9,OLFM4), Prognosis(SM3,desmin,surviving,hTERT and NM23), Potential therapeutic targets(EB1)[22-23]. Therefore, exploring the protein interaction network of tumors, analysing the characteristics of the protein molecular interaction network, and excavating a variety of signalling pathway and the genes will provide a biological basis for the study of the molecular mechanism of tumor and the further treatment as well.

Bio-molecular network analysis is an important direction of systematic biology research. Large-scale human protein-protein network can provide new insights into protein functions, pathways, molecular machines and functional protein modules [24]. The function of bio-molecular often depends on modularization, the network module is made by a number of nodes in the conjunction of each other and has a stable structure which can often reflect a similar nature between the nodes [25, 26]. Analysing the function module is the one of the most common method to analyse biological molecular network. According to 127 genes provided by OMIM, our research has built up protein interaction network of rectal cancer, which contain 996 nodes (proteins), 3377 edges (interaction). Due to the network is very large, the experimental introduced MCOMD algorithm to evaluate the network’s regional integration through the correlation integral. Correlation integral descripted the proteins associated with the degree in the region. Proteins of the same molecular complex generally have the same biological function, therefore we can discover the unknown gene functions or new molecular functional groups, such as cluster 1.

Gene *DDK1, sparcl1, wisp2, cux1, pabpc1, ptk2* and *htra1* as the composition members of the cluster1(score 20) in the centre, have closely relationships with each other as well as other genes. It has reported that the secreted protein acidic and rich in cysteine-like 1 *(,sparcl1)* is expressed in various normal tissues and many types of cancers. Another study has shown that marker *sparcl1* was significantly related to the prognosis and clinical pathological features of the CRC patients [27]. *Sparcl1* expression increased with RT and is related to a better prognosis in rectal cancer patients with RT but not in patients without RT. This result may help us to select the patients to the best suited preoperative RT. In the process of rectal cancer, gene WISP2 knockout significantly increased Caco-2 cell invasion and motility. Up-regulation of MMP2, -7 and -9 may indicate that WISP2regulates invasion and motility through MMPs. Regulation of invasion by WISP2 may involve the WNT signalling pathway[28]. Thus, some studies have identified CUX1 as a pan-driver of tumorigenesis and uncover a potential strategy for treating CUX1-mutant tumors. So from cluster1 we predict the effect of gene Cut1 to rectal cancer may be that CUX1 deficiency activates phosphoinositide 3-kinase (PI3K) signalling through direct transcriptional downregulation of the PI3K inhibitor PIK3IP1 (phosphoinositide-3-kinase interacting protein 1), leading to increased tumor growth and susceptibility to PI3K-AKT inhibition. And we also can forecast a few clones existing in colorectal cancer, containing gene mutation of ptk2, htra1 and PABPC1.

Rectal cancer is demonstrated not simply controlled by a particular gene or signaling pathways, but also by the complex process of network system co-ordinately regulated which consisted of a variety of signalling pathways and multiple genes. In the signalling network, it is likely there is some " key regulatory point". At last, a best understanding of chemotherapy molecular targets allowed the identification of genetic markers that can predict the response and/or the toxicity of anti-cancer drugs used in rectal cancers, which could be helpful in the future to propose for each patient a personalized treatment [29-31]. Mutations that can predict the response of new target therapies such as the inhibitors of the JAK kinase inhibitor AG4 90 in colorectal cancer have also been found and will allow the selection of patients who can have benefit from these new therapeutic drugs. The experiment dig out a variety of signalling pathways and genes which can provide reliable directions for molecular mechanism research of treatment, and it need to be further verified.

### Compliance with Ethical Standards

**Conflict of interest:** We declare that we have no financial and personal relationships with other people or organizations that can inappropriately influence our work, there is no professional or other personal interest of any nature or kind in any product, service and/or company that could be construed as influencing the position presented in, or the review of, the manuscript entitled exploring genes of rectal cancer for new treatments based on protein interaction network

**Ethical approval:** This article does not contain any studies with human participants or animals performed by any of the authors.

